# A Deep Recurrent Neural Network Discovers Complex Biological Rules to Decipher RNA Protein-Coding Potential

**DOI:** 10.1101/200758

**Authors:** Steven T. Hill, Rachael Kuintzle, Amy Teegarden, Erich Merrill, Padideh Danaee, David A. Hendrix

## Abstract

The current deluge of newly identified RNA transcripts presents a singular opportunity for improved assessment of coding potential, a cornerstone of genome annotation, and for machine-driven discovery of biological knowledge. While traditional, feature-based methods for RNA classification are limited by current scientific knowledge, deep learning methods can independently discover complex biological rules in the data *de novo*. We trained a gated recurrent neural network (RNN) on human messenger RNA (mRNA) and long noncoding RNA (lncRNA) sequences. Our model, mRNA RNN (mRNN), surpasses state-of-the-art methods at predicting protein-coding potential. To understand what mRNN learned, we probed the network and uncovered several context-sensitive codons highly predictive of coding potential. Our results suggest that gated RNNs can learn complex and long-range patterns in full-length human transcripts, making them ideal for performing a wide range of difficult classification tasks and, most importantly, for harvesting new biological insights from the rising flood of sequencing data.

## Introduction

Deep sequencing technology has yielded a torrent of new transcript annotations, creating a need for fresh approaches to unlock the full information potential of these vast datasets. Existing state-of-the-art methods for classification of long RNAs as protein-coding RNAs (mRNAs) or long noncoding RNAs (lncRNAs) rely on human-engineered features, such as the coverage and length of a predicted open reading frame (ORF). These features predispose such models to misclassification of mRNAs encoding small proteins and of lncRNAs with long, un-translated ORFs. Nucleotide hexamer frequency is another commonly used feature, but while it can capture the frequency of codon pairs, it does not benefit from the larger sequence context. These limitations and the annotation challenges ahead demand new approaches to biological sequence classification that are capable of detecting complex, variable-length patterns.

In contrast to conventional machine learning methods, “deep learning”–the application of multi-layered artificial neural networks to learning tasks–can discover useful features independently, avoiding biases introduced by human-engineered features(1). Deep learning methods have repeatedly outperformed state-of-the-art “shallow” machine learning algorithms, such as support vector machines (SVM) and logistic regression, as approaches to biological problems in recent years. Multiple bioinformatics applications of deep convolutional neural networks (CNNs) have been published(2-4); however, while CNNs adeptly learn spatial information, recurrent neural networks (RNNs) are better suited for learning sequential patterns because of their serialized structure and ability to handle variable-length inputs(1). Following the success of RNN-based approaches in the fields of natural language and music(5), researchers have only recently begun to apply RNNs to biological problems(6-10). While basic RNNs are challenged by most biologically-relevant input sequence lengths due to the “vanishing gradient problem,” a difficulty encountered during training due to the multiplication of many small terms when computing the gradient of an error function by the chain rule (11), several recent adaptations addressed this issue. Among the most popular of these modified RNNs are long-short-term-memory (LSTM) RNNs (12) and gated recurrent unit (GRU) RNNs (13), which manage memory to improve the learning of long-range dependencies. Recent studies demonstrated superior performance of GRUs compared to LSTMs for bioinformatics tasks(7,14). We report the successful implementation of a GRU network to accurately predict protein-coding potential of complete, variable-length transcripts (Fig. 1). Our method, “mRNA RNN” (mRNN), not only outperforms existing state-of-the-art classifiers, but also learns complex biological rules in the process.

**Figure 1.**
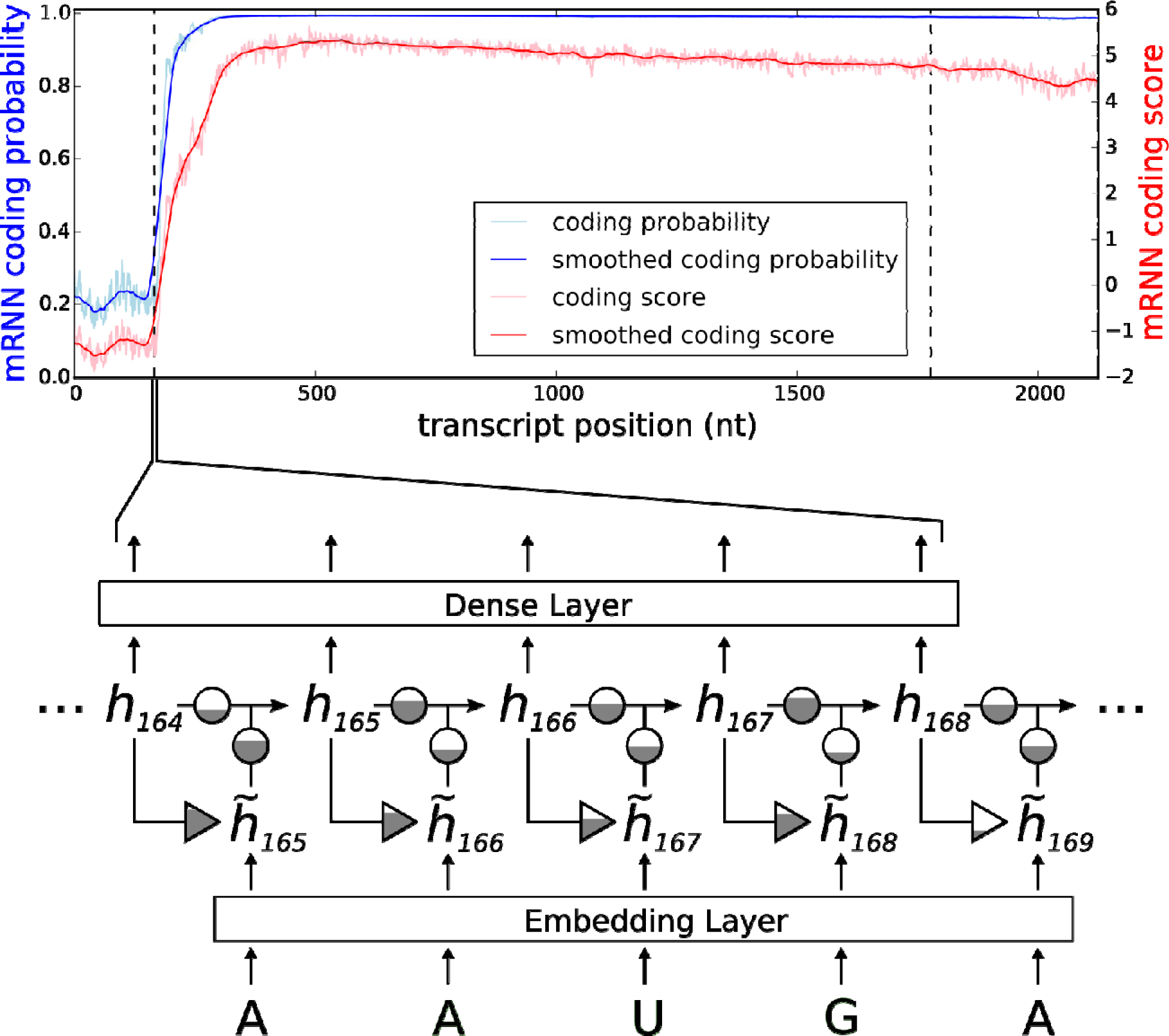
mRNN Output and Model Schematic. Coding probability and coding potential score is shown at nucleotide-level resolution for the transcript ENST00000371732.9, which encodes caspase recruitment domain family member 9. Values at position correspond to the mRNN coding probability or, the mRNN output for the truncated sequence from 1 to . Vertical dashed lines demarcate the annotated start and end of the CDS. A schematic of the gated recurrent neural network is shown below. Equilateral triangles signify reset gates, and the height of the grey fill represents the proportional contribution of the previous hidden state ( ) to the new candidate hidden state ( ). The update gate is shown as two circles representing the proportional contributions of the previous hidden state ( ) and the new candidate hidden state ( ) to the new hidden state ( ). Arrow represent matrix products. The embedding layer maps nucleotides to 128-dimnensional vectors.

## Results

The best resulting mRNN model after training selected by accuracy on the validation set is referred to hereafter as “mRNN”. We also implemented an ensemble testing method called “mRNN ensemble,” which uses the weighted average of the five best mRNN models. For comparison, we used the same test set to assess performance of two non-comparative classifiers considered to be state-of-the-art in speed and performance: CPAT(15) and FEELnc(16). While mRNN matched or outperformed CPAT and FEELnc on the test set, the mRNN ensemble method showed significant improvements in performance over these methods in accuracy, specificity, and other metrics (Fig. 2A). We also compared the classifiers using a set of more challenging transcripts, including 500 mRNAs with short ORFs (≤ 50 codons), and 500 lncRNAs with long (untranslated) ORFs (≥ 50 codons) (Fig. 2B). Both mRNN and mRNN ensemble methods significantly outperformed CPAT and FEELnc in all metrics on this challenge set. Notably, CPAT and FEELnc showed low sensitivity (62.2% and 67.8% respectively), indicating a bias against classification of mRNAs with short ORFs as protein-coding, while mRNN ensemble achieved a sensitivity of 78.6%, demonstrating its superior predictive power for these atypical transcripts.

**Figure 2.**
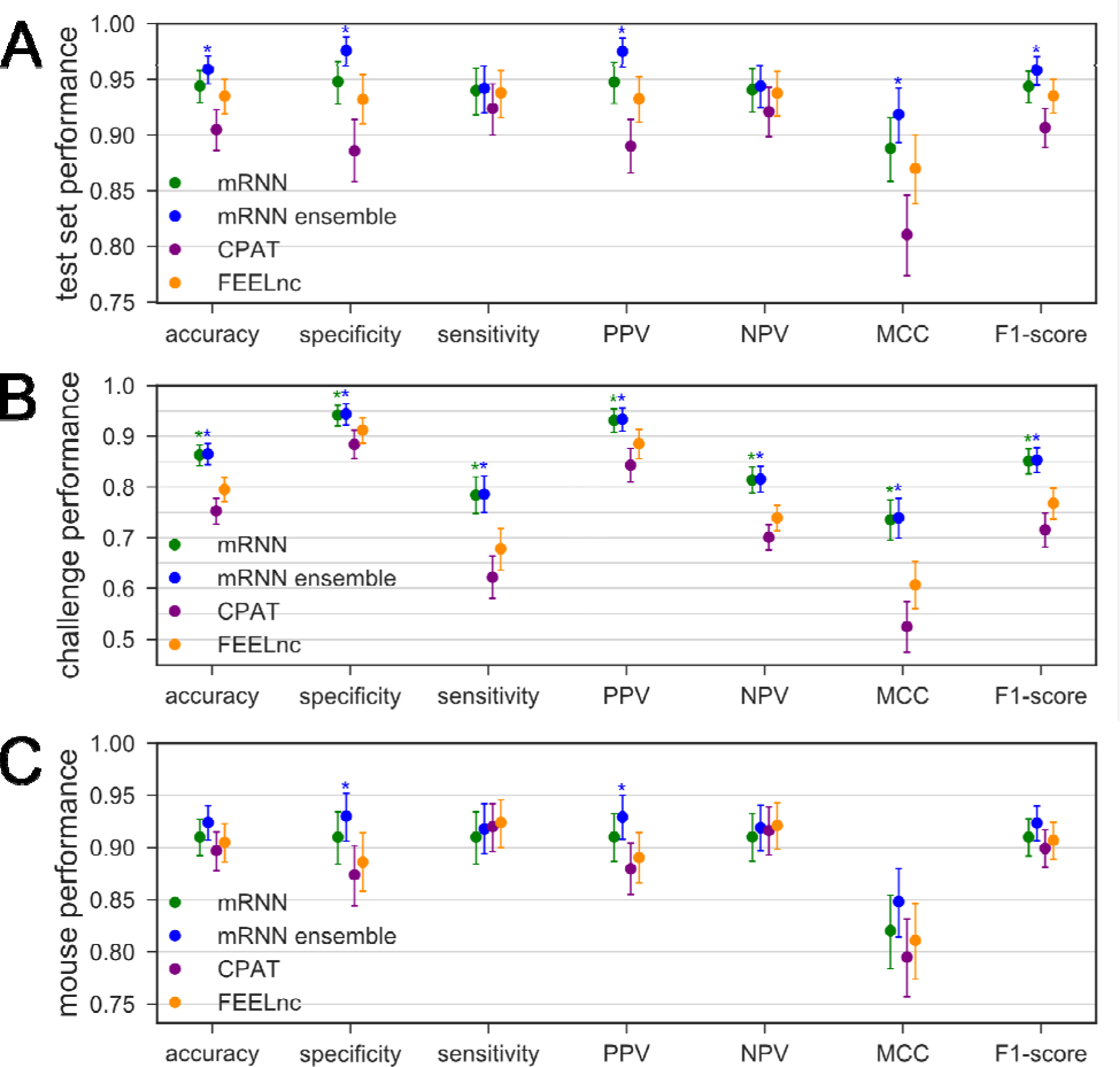
Comparison of Classifier Performance. (A-C) Performance of four classifiers trained with human transcript sequences. Error bars are 95% confidence intervals computed from 100,000 bootstrap trials. Asterisks above mRNN or ensemble mRNN indicate the method's improvement over both CPAT and FEELnc with an empirical p-value less than 0.05 computed from the bootstrap trials. (A) Performance on human test set transcripts, consisting of 500 mRNAs and 500 lncRNAs. (B) Performance on human challenge set transcripts, including 500 mRNAs with ORFs<50 codons and 500 lncRNAs with ORFs>50 codons. (C) Performance on a test set of mouse transcripts, including 500 mRNA and 500 lncRNAs, using models trained with human data.

We next evaluated all models on a randomly selected test set from mouse GENCODE transcripts consisting of 500 mRNAs and 500 lncRNAs. For each method, we trained on the same human training set, and tested on mouse. The mRNN ensemble and mRNN classifiers had the highest accuracy and specificity, showing that they can be used for classification of long RNAs in a new transcriptome (Fig. 2C).

To begin deducing what mRNN learned, we conducted sequence perturbation analyses (Supplementary Methods). Score changes for sequences with shuffled coding sequence (CDS) regions compared to those with shuffled 3’ or 5’ untranslated regions (UTRs) demonstrate that mRNN primarily utilizes organized sequence information in the CDS (Fig. S6). We next conducted a point-mutation analysis to evaluate changes in score resulting from every possible single-nucleotide substitution for all GENCODE mRNA transcripts under 2,000 nt in length (Fig. 3). The annotated start codon marked a clear boundary, with low score changes preceding it and strong changes following it, indicating that sequence perturbations early in the CDS erase more predictive information than perturbations in the 5’ UTR. In contrast, score changes around non-start AUGs in the 5’ UTR were more symmetric before and after the AUG, and significantly lower on average. Strikingly, the pattern of average score changes in the true CDS exhibited three-nucleotide periodicity with a persistent aversion to mutations that made codons more similar to in-frame stop codons. This pattern was not observed upstream of the annotated start codon (5’ UTR), nor in the regions flanking either AUGs in the 5’ UTR or control CUGs (Fig. S7). An aversion to stop codon-like trinucleotides was also observed preceding, but not following, annotated stop codons, suggesting that mRNN recognizes the end of the CDS. This pattern was not observed in regions preceding UGA/UAA/UAG trinucleotides in the 3’ UTR. Notably, mutation of the annotated stop codon significantly increased the coding potential score, showing that mRNN displays a preference for longer ORFs.

**Figure 3.**
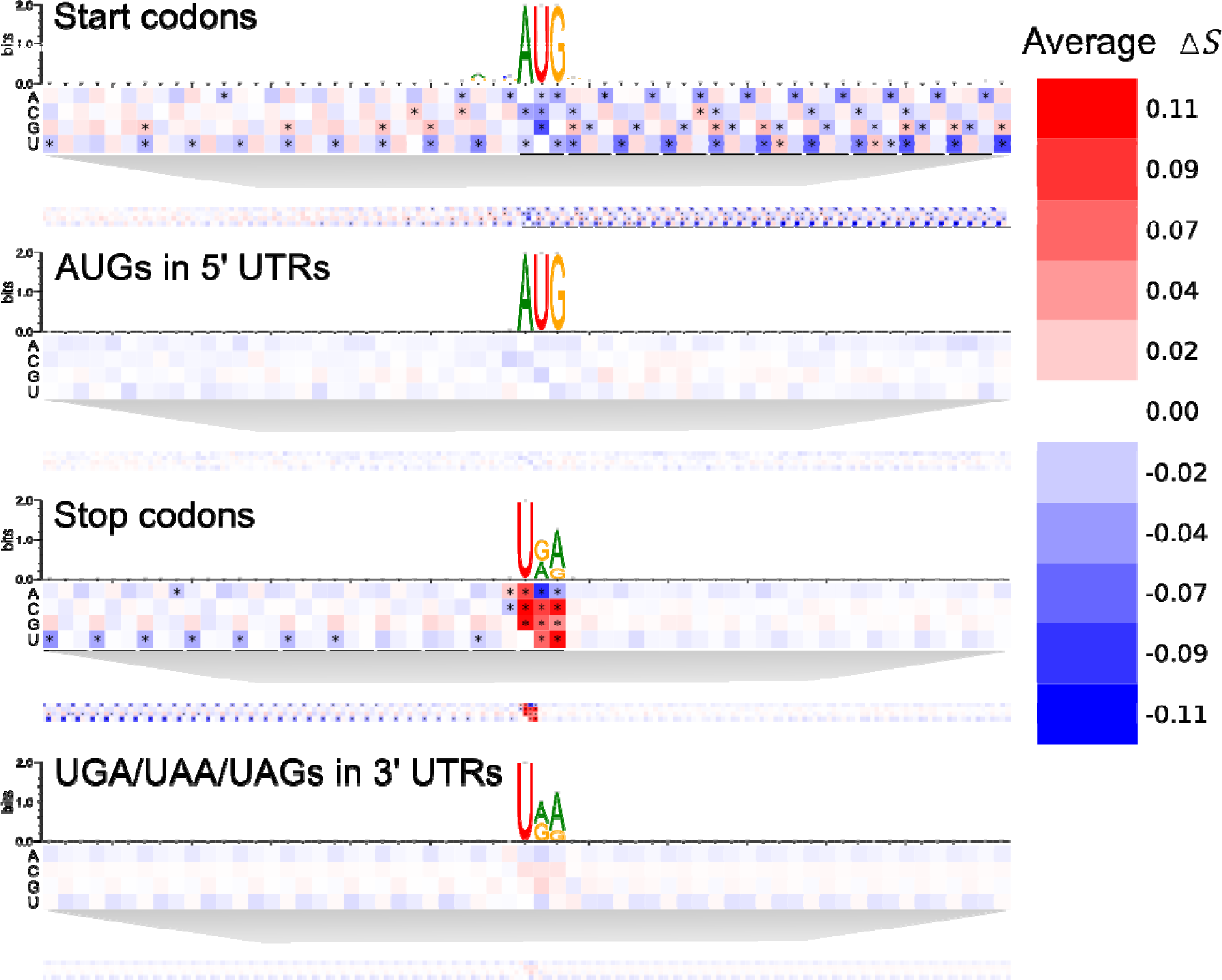
Transcript Point Mutation Maps. Heat maps representing the average change in score for point mutations at positions relative to the following elements (from top to bottom): annotated start codons, AUGs in 5’ UTRs, annotated stop codons, and UGA/UAA/UAGs in 3’ UTRs. Sequence logos present the nucleotide composition of the sequences analyzed around the same windows. Asterisks represent cells that are statistically significant at an FDR of 0.0001 using a two-tailed t-test comparing score changes from mutations at a given position to all score changes from mutations of the same base in the corresponding background region. Background regions are 5’ UTRs for the start codons and AUGs, or 3’ UTRs for the stop codons and UGA/UAA/UAGs.

To evaluate whether mRNN learned relationships between distinct features, we performed pairwise-mutations of the transcript SPANXB1, a cancer/testis-specific antigen. This transcript is relatively short (under 500 nt) and has both 5’ and 3’ UTRs. The coding score for this transcript starts an upward trajectory within the CDS (Fig. 4A). We examined this transcript by altering every possible combination of two nucleotides within the sequence. We define Δ*S_syn_(i, j)* as the minimum of the difference between the coding score change resulting from mutations at two positions i and j and the sum of score changes associated with the individual mutations. Therefore, Δ*S_syn_(i, j)* quantifies the “score change synergy” of the pair of mutations, and is strongly negative for highly related positions. We identified several pairs of synergistic mutations, including a point mutation that changed an AAC codon to AAA and another that changed an AAG codon to UAG, which, when combined, resulted in a score change 6.8 times the sum of the individual score changes (Fig. 4B). In other examples, we identified compensatory changes, such as a decrease in score from a nonsense mutation that was significantly diminished when another mutation changed a UGU codon to UAU (Fig. 4C). In both cases described, the two mutated positions are located within the coding score spike, despite being separated by 18 and 29 bases, respectively. The fact that some mutations can exacerbate or compensate for the effect of ORF-truncating mutations on mRNN's output demonstrates that mRNN learned complex and long-range sequence information dependencies.

**Figure 4.**
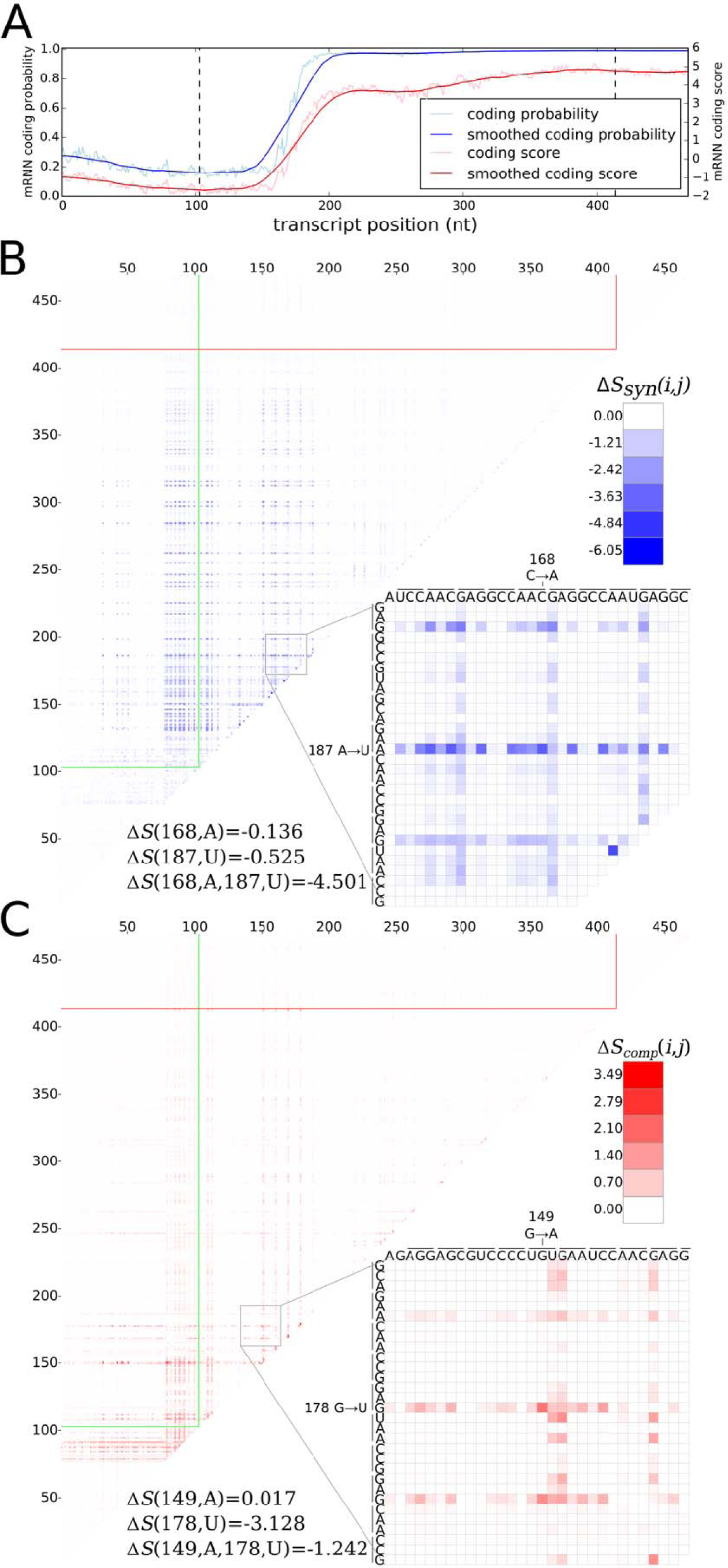
Pair-wise mutation analysis. (A) The mRNN coding trajectory (as in Fig. 1), for ENST00000449283.1, a transcript encoding SPANX1. (B) Pair-wise mutation heat map of synergistic score changes for the same transcript. Values are the score change synergy for a pair of mutated bases at positions *i* and *j*, where *i < j*. Score change synergy is the minimum difference between the resulting change in score when the pair of bases is mutated and the sum of the score changes from individual mutations of each base in the pair. (C) Pair-wise mutation heat map of compensatory score changes for the same transcript. Values are the compensatory score change for a pair of mutated bases at positions *i* and *j*, where *i < j*. Compensatory score change is the maximum difference between the resulting change in score when the pair of bases is mutated and the sum of the score changes from individual mutations of each base in the pair. (B-C) Bottom-right of each heat map shows a zoomed-in view of a position pair with a highly compensatory or synergistic score change. Each line spanning three nucleotides represents a codon.

To visualize mRNN's decision-making process, we defined a coding trajectory *S_trunc_(i)* as the score of the truncated sequence from 1 to *i* for transcripts in the human test set (Fig. 5A). Remarkably, we found several examples of mRNAs with long 5’ UTRs that mRNN classified as coding only after observing the CDS more than 4,000 nt from the transcript start (Fig. S8A-D). Thus, mRNN remains sensitive to information toward the end of transcripts longer than sequences previously used in any bioinformatics RNN applications that we are aware of, despite being trained only on sequences shorter than 1,000 nucleotides long.

**Figure 5.**
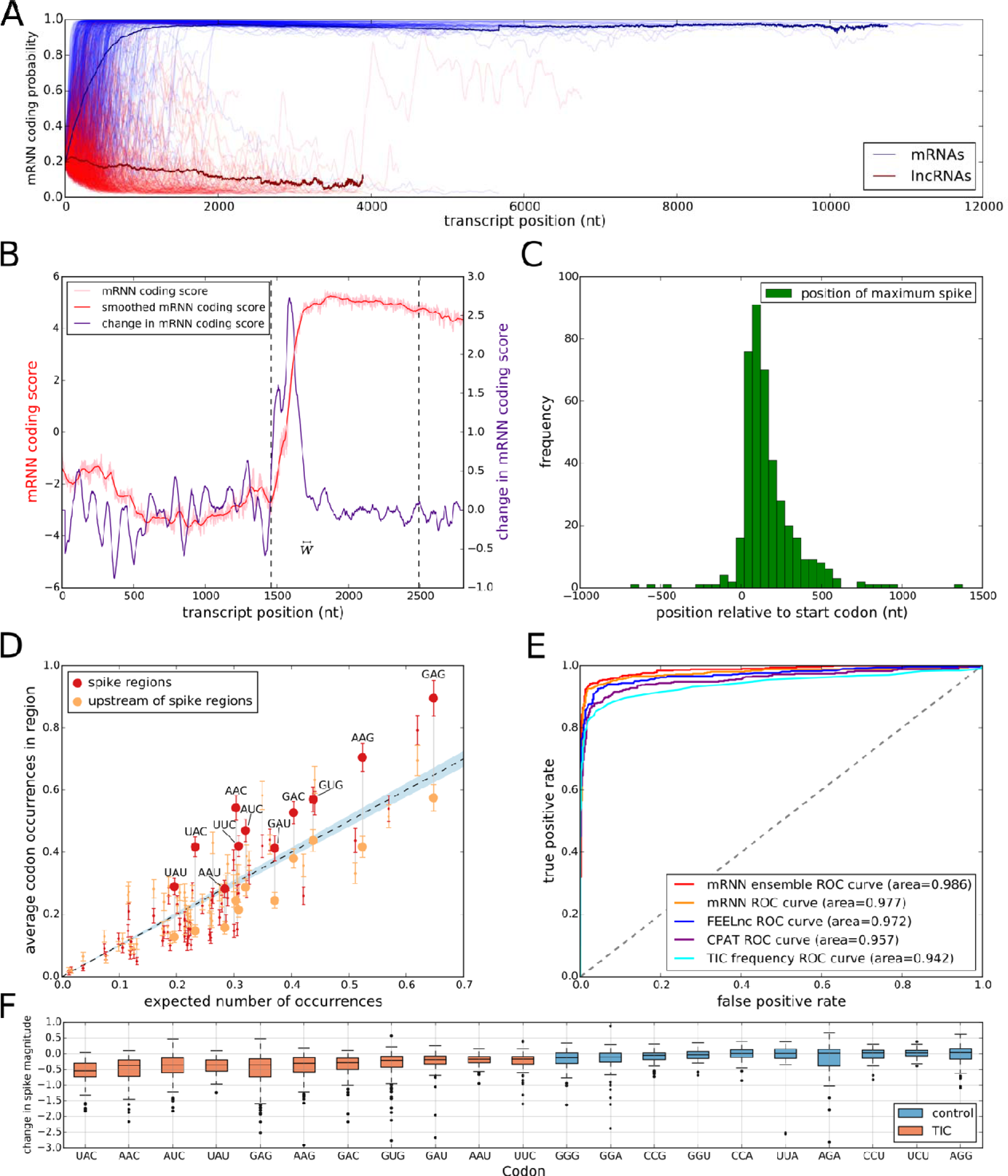
Model Interrogation for Feature Discovery. (A) mRNN coding score trajectories without smoothing for each transcript in the test set. Blue, protein-coding; red, noncoding. Bold lines represent average coding probability when 5 or more transcripts had lengths at of least nt. (B) Coding score trajectory for transcript ENST00000458629.1, which encodes C-X-C motif chemokine receptor 6. Vertical dashed line mark CDS boundaries. (C) Histogram of significant spike locations in test set mRNAs relative to true CDS start positions. (D) Scatterplot showing codons enriched in the spike regions (+/− 25 nt around most significant spike position) compared to 50-nt regions upstream of the spikes. The x-axis is the frequency of each codon in the full set of GENCODE annotated coding regions. The y-axis represents the frequency of the codon in the indicated region. Each pair of points represents a codon. Large, labeled points are translation-indicating codons (TICs)—codons statistically enriched (FDR 0.05) in spike regions compared to the regions upstream of spikes. The dashed line corresponds to global codon frequency, and the blue band is the range of standard error computed from a binomial model. (E) Receiver operator characteristic analysis for five prediction methods including our mRNN ensemble, the best single mRNN model, FEELnc, CPAT, and TIC frequency. TIC frequency is the number of occurrences of TICs within 1000 nt of, and in-frame with, an upstream AUG, but not after an in-frame UGA/UAA/UAG. AUROC values for each method are presented in the legend. (F) mRNN coding score changes resulting from *in silico* TIC mutations. While the majority of mutations to TICs lead to a decrease in coding score, mutations to control codons (the codons least enriched in the spike regions) result in smaller score changes on average.

To identify regions of the sequence that most strongly impact mRNN's decision, we performed unweighted sliding-average smoothing of the coding potential trajectories, then computed the change in score *ΔS_trunc_(i)* across the sequence for a window w of 50 nucleotides (Fig. 5B). Statistically significant spikes (Fig. S9) were identified in 412 of the 500 test mRNAs, and in only 47 of the 500 lncRNAs. The distribution of the spike positions for mRNAs peaked within the CDS, shortly after the start codon (Fig. 5C, Fig. S10, and S11A-B).

To identify the sequence elements associated with significant spikes in coding potential score, we compared the frequency of in-frame codons in a 50-nt window centered at the spike to codon frequencies in the 50-nt window preceding the spike. We found 11 significantly enriched codons using a t-test and an FDR of 0.05 (Fig. 5D, Table S1); we named these translation-indicating codons (TICs). 9 of the 11 TICs were also significantly enriched in spike regions of an independent set of mRNAs with long 5’ UTRs (Fig. S11C). Notably, two codons in the synergistic and compensatory pairwise mutation examples above (AAC and UAU) are TICs.

To assess the predictive power of TICs, we defined a TIC-score as the maximum number of TICs occurring within 1000 nt downstream of an in-frame AUG, and preceding the first in-frame stop codon. This TIC-score was able to accurately predict coding potential in the test set with an AUROC of 0.942, just below that of CPAT at 0.957 (Fig. 5E). The same rule distinguished mRNAs from lncRNAs in the full GENCODE datasets with an AUROC of 0.931 for human and 0.935 for mouse. We next computed the reduction in the spike magnitude—the change in *ΔS_trunc_(i)*—resulting from the mutation of a given TIC codon *in silico*. Mutation of TICs resulted in spike height decreases 94.7% of the time, while mutations to the least enriched codons in the spike regions decreased spike height only 59.9% of the time (Fig. 5F, S12), demonstrating that TICs are an important part of mRNN's classification process. Of note, we also identified a frame-biased, 12-mer motif enriched in spike regions, but it possessed lower predictive power than the TICs (Fig. S13, Table S2-3). Strikingly, the TICs are among the codons most enriched in GENCODE CDSs relative to UTRs and out-of-frame triplets, demonstrating that mRNN learned the complex sequence context that gives these codons predictive power (Fig. S14).

## Discussion

In this study, we have shown that GRU networks can successfully model full-length human transcripts. Previous bioinformatics applications of RNNs restricted input sequence length to 2,000 nt or fewer by one of three strategies: filtering the dataset on a length threshold(17), dividing input sequences into segments of a fixed size(18,19), or truncating input sequences(20). However, one important advantage that deep RNNs have over other deep learning methods is the ability to interpret context and long-range information dependencies. In order to exploit the full power of our GRU network, we did not truncate or segment our training sequences, and we did not constrain our test set inputs by sequence length in any way. Our test inputs included 222 mRNAs longer than 2,000 nucleotides, and our model showed no impairment in classifying even the longest sequences, which exceeded 100,000 nucleotides.

Despite mRNN's featureless architecture, which precluded it from integrating human knowledge of mRNA structure into its learning process, mRNN was able to learn true defining features of mRNAs, including trinucleotide patterns and depletion of in-frame stop codons after the start of an open-reading frame. Furthermore, the recurrent nature of mRNN enabled it to leverage long-range information dependencies for classification, as evidenced by pairwise mutation analysis. In addition to surpassing state-of-the-art accuracy in assessment of transcript coding potential, we demonstrate that the “black-box” GRU network can be harnessed for identifying specific biological attributes, such as the translation-indicating codons (TICs), that distinguish sequence classes. We anticipate that GRU-based approaches will be highly useful for future bioinformatics classification tasks, as well as for unlocking hidden biological insights in the vast amounts of available sequencing data.

## Materials and Methods

For training, we provided mRNN with a dataset containing full-length human transcript sequences labeled as mRNAs or lncRNAs. All training and test sets were selected from GENCODE Release 25(21). We evaluated mRNN's performance using a test set—an unbiased random sample of human transcripts composed of 500 mRNAs and 500 lncRNAs selected from the full GENCODE annotation. Similarly, we selected a validation set for hyper-parameter tuning and model selection (Fig. S1-3) equal in size to the test set. No length limit or other filter was imposed on sequences in the test or validation sets.

To reduce computation time and to prevent the learning of transcript length as a feature, we imposed constraints on the training set sequence length for mRNN. After selecting the test and validation set transcripts, we selected 16,000 mRNAs and 16,000 lncRNAs from the remaining sequences between 200 and 1000 nt long as our training set. We used a combination of “data augmentation,” in which we pre-train models on mutated copies of the training set, and “early stopping,” which exits training if loss on the training set decreases while validation loss does not; both of these strategies help prevent over-fitting during training. Model parameters were selected based on validation loss during hyper-parameter tuning. We used embedding vectors to represent each nucleotide because this yielded higher validation accuracy than did one-hot encoding when using ensemble testing for the RNN library, Passage. We also used dropout, which randomly sets network inputs to zero in Passage's GRU implementation, because it improved validation accuracy with embedding. No length restrictions were imposed on sequences in the training set for CPAT and FEELnc, giving these classifiers substantially more training data than mRNN. For detailed methods see Supplementary Methods.

## Software availability

Source code implementing data preprocessing, training, and downstream analysis is available in the package mRNN from http://github.com/hendrixlab/mRNN.

## Acknowledgements

The authors would like to thank Prof. Stephen Ramsey, Prof. Christopher K. Mathews, Prof. Liang Huang, Prof. Colin Johnson, Prof. P. Andy Karplus, and Prof. Michael Freitag for feedback on the manuscript and helpful discussions.

